# Development of the First Tractable Genetic System for *Parvimonas micra*, a Ubiquitous Pathobiont in Human Dysbiotic Disease

**DOI:** 10.1101/2022.01.27.477930

**Authors:** Dustin L. Higashi, Sean McGuire, Yasser Abdelrahman, Zhengzhong Zou, Hua Qin, David Anderson, Elizabeth A. Palmer, Jens Kreth, Justin Merritt

## Abstract

*Parvimonas micra* is a Gram-positive obligate anaerobe and a typical member of the human microbiome. *P. micra* is among the most highly enriched species at numerous sites of mucosal dysbiotic disease and is closely associated with the development of multiple types of malignant tumors. Despite its strong association with disease, surprisingly little is known about *P. micra* pathobiology, which is directly attributable to its longstanding genetic intractability. To address this problem, we directly isolated a collection of *P. micra* strains from odontogenic abscess clinical specimens and then screened these isolates for natural competence. Amazingly, all of the *P. micra* clinical isolates exhibited various levels of natural competence, including the reference strain ATCC 33270. By exploiting this ability, we were able to employ cloning-independent methodologies to engineer and complement a variety of targeted chromosomal genetic mutations directly within low passage clinical isolates. To create the first *P. micra* genetic system, we employed renilla-based bioluminescence for highly sensitive reporter studies. This reporter system was then applied for the development of the novel Theo+ theophylline-inducible riboswitch for tunable gene expression studies over a broad dynamic range. Finally, we demonstrate the feasibility of generating Mariner-based Tn-seq libraries for forward genetic screening in *P. micra*. With the availability of a highly efficient transformation protocol and the current suite of genetic tools, *P. micra* should now be considered as a fully genetically tractable organism suitable for molecular genetic research. The methods presented here provide a clear path to investigate the understudied role of *P. micra* in polymicrobial infections and tumorigenesis.

## Introduction

*Parvimonas micra* is a Gram-positive obligate anaerobe from the largely uncharacterized Tissierellia class of the Firmicutes phylum. Initially grouped within the *Peptostreptococcus* genus, a taxonomic reclassification revealed its distinction within the Firmicutes [1, 2]. *P. micra* is a common commensal of microbiomes at various mucosal sites in the body, including the oral cavity, gastrointestinal tract, respiratory system, and female urogenital tract. *P. micra* is a surprisingly common and abundant species detected in numerous epidemiological studies of different mucosal inflammatory diseases [3]. In addition, *P. micra* is among the most common sources of Gram-positive anaerobic cocci (GPAC) sepsis. It is a major constituent of numerous types of systemic abscesses, and it exhibits a strong association with a variety of malignant tumors [4–7]. Indeed, its presence has even been proposed as a discriminating biomarker for colorectal cancer, gastric cancer, and oral cancer [4, 8, 9]. In the oral cavity, *P. micra* is highly enriched in periodontitis lesions, infected root canals, and is especially prevalent in polymicrobial odontogenic abscesses [10–15].

The mechanisms by which *P. micra* contributes to human health and disease remain largely enigmatic, as there is a severe paucity of literature describing its molecular genetics [7]. Furthermore, this organism has been historically challenging to identify in clinical microbiology laboratories, largely due to its fastidious nature and slow growth rate [3]. Reports suggest that *P. micra* frequently serves as a major pathobiont involved in pathogenic synergism with other members of the microbiome. A recent *in vitro* study illustrated how *P. micra* can augment the growth of *Porphyromonas gingivalis* and enhance its production of secreted proteolytic gingipains [16]. *In vivo* studies demonstrated an enhanced transmissibility of pus generated from *P. micra/Prevotella* co-infections compared to their respective mono-infections [17]. *P. micra* can also coaggregate with the oral pathobionts *Treponema denticola* and *Fusobacterium nucleatum*, implicating a potential role in polymicrobial oral biofilm formation [18, 19].

Despite its strong association with a broad diversity of mucosal inflammatory diseases, *P. micra* remains vastly understudied, largely due to its genetic intractability. Only a single report of targeted mutagenesis has ever been described for this organism and this was accomplished via electroporation of a suicide vector [20]. The lack of follow-up genetic studies underscores the significant challenges associated with genetically manipulating *P. micra*. Such problems have remained a significant deterrent for detailed mechanistic studies of *P. micra* pathobiology. To address this problem, we present the first efficacious *P. micra* genetic system, which is founded upon a newly discovered natural competence ability in this organism. By exploiting this ability, we were able to apply cloning-independent methodologies to engineer a variety of targeted chromosomal genetic modifications using low passage clinical isolates. The efficacy of our *P. micra* transformation protocol further supported the development of the first tractable genetic system for this species. With the genetic toolbox presented here, a complete molecular genetic system is in place to reliably interrogate *P. micra* pathobiology.

## Results

### Identification of Natural Competence in *P. micra* clinical isolates

Recently, Liu and Hou described the only known report of targeted mutagenesis in *P. micra* [20]. This study employed a cloning-based methodology to create a mutagenesis construct introduced via electrotransformation of a suicide plasmid. Due to the difficulties associated with *P. micra* electroporation, we were interested to determine whether a more reliable transformation approach could be developed using natural competence. To test this, we first isolated a collection of low passage clinical strains directly from odontogenic abscess specimens (Figure 1A). Following tooth extraction, abscess pus samples were collected into pre-reduced transport media [21] and then spread onto PMM selective/differential agar media [12]. Candidate *P. micra* colonies were selected based upon the accumulation of black precipitate, then examined morphologically by microscopy, and finally confirmed with 16S rRNA sequencing (Figure 1A).

**Figure 1.**
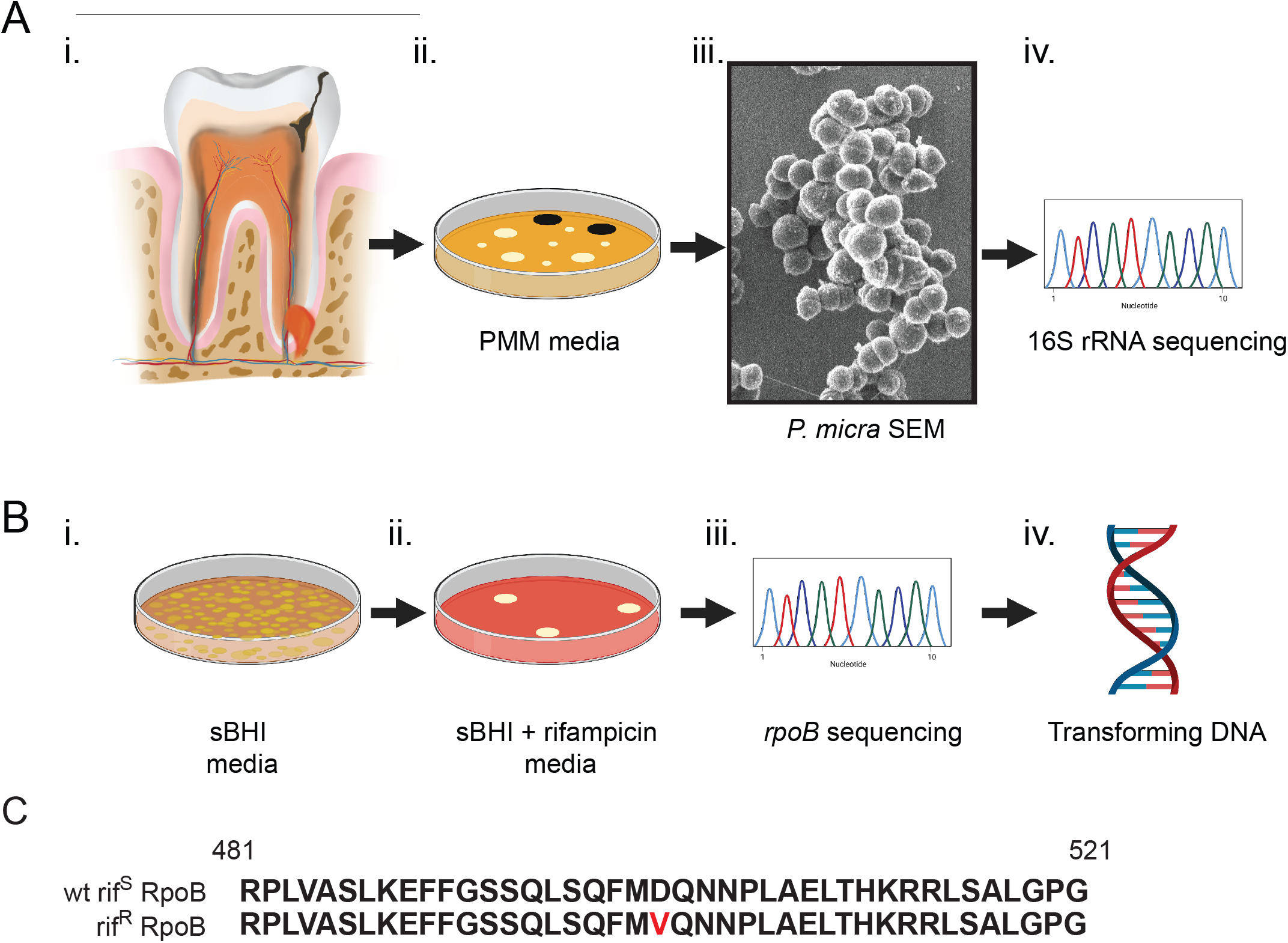
*Parvimonas micra* clinical isolation scheme. (A) (i) Odontogenic abscess samples were collected into anaerobic transport media and (ii) grown on PMM selective/differential agar. Putative colonies were (iii) confirmed morphologically and (iv) submitted for 16S rRNA sequencing to verify *P. micra*. (B) Wild-type strain ATCC 33270 was (i) grown on non-selective sBHI agar and (ii) subsequently placed on rifampicin containing sBHI media to isolate a spontaneous rifampicin resistance strain (rif^R^). (iii) The *rpoB* gene was sequenced and (iv) genomic DNA was isolated from both the rif^R^ and wild type (wt) rifampicin sensitive (rif^S^) strain. (C) The deduced amino acid sequences of the wild-type rif^S^ and rif^R^ RpoB proteins are shown. The rif^R^ mutation is shown in red font.

We next generated a selectable marker for use in DNA transformation assays by isolating a spontaneous rifampicin resistant mutant of the wild-type reference strain ATCC 33270 (Figure 1B). A resulting rifampicin resistant (rif^R^) colony of ATCC 33270 was verified by sequence analysis of the *rpoB* gene encoding the beta subunit of RNA polymerase. A point mutation in this gene confirmed a D501V substitution in RpoB (Figure 1C), which coincides with rifampicin resistance mutations detected in other bacterial species [22, 23]. Next, genomic DNA (gDNA) was extracted from both the wild-type strain ATCC 33270 (rif^S^) and the rifampicin resistant mutant of ATCC 33270 (rif^R^) (Figure 1B) and then transformed into our collection of wild-type *P. micra* clinical isolates using a transformation protocol similar to that previously employed for *Veillonella parvula* [24]. With this approach, we observed natural competence from the majority of strains tested, with particularly high transformation efficiencies in a number of isolates (Table 1). These results confirmed that *P. micra* is capable of developing natural competence.

**Table 1.** Transformation frequency is expressed as the number of rifampcin resistant bacteria/ total CFUs. Values are averaged from triplicate determinations ± SD. Statistical analysis was performed using a two-tailed Student’s *t*-test.

| *Parvimonas micra*<br>Isolate | ATCC<br>33270<br>gDNA<br>(rif<sup>R</sup>) | Transformation<br>Frequency (rif<sup>R</sup>/total<br>CFUs)<br>(x 10<sup>-4</sup>) | Fold-increase<br>(over<br>background) |
| --- | --- | --- | --- |
| A28 | + | 1.73 ± 0.34 * | 752 |
| A28 | - | 0.0023 ± 0.0005 |  |
| A42 | + | 0.33 ± 0.12 * | 236 |
| A42 | - | 0.0014 ± 0.0002 |  |
| A11 | + | 0.26 ± 0.06 * | 263 |
| A11 | - | 0.00099 ± 0.00014 |  |
| ATCC 33270 | + | 0.075 ± 0.0029 * | 36 |
| ATCC 33270 | - | 0.0021 ± 0.0022 |  |
| A3 | + | 0.013 ± 0.00013 * | 8 |
| A3 | - | 0.0016 ± 0.00043 |  |
| A1 | + | 0.0040 ± 0.0012 <sup>n.s.</sup> | 1 |
| A1 | - | 0.0035 ± 0.0014 |  |

“\*” indicates significance ( $p < 0.05$ ) relative to wt gDNA control. “n.s.” indicates no significance ( $p > 0.60$ ).

### Cloning-Independent Targeted Mutagenesis

Following our discovery of *P. micra* natural competence, it was of interest to determine whether this ability could be exploited for engineering targeted mutations in *P. micra*. For this, we chose the *ermB* erythromycin resistance cassette as a selectable marker [25]. An allelic replacement mutagenesis construct was created by Gibson assembly of PCR amplicons containing *ermB* flanked by one kb homologous fragments of the EF-Tu-encoding gene *tuf* and its downstream locus (Figure 2A, top). Transformation of this construct into the clinical strain A28 resulted in the insertion of *ermB* immediately downstream of the *tuf* open reading frame (ORF) through a double crossover allelic exchange event (Figures 2A and B). Next, replicate constructs were generated for each isolate and then transformed into their respective strains, including the reference strain ATCC 33270. In each of these reactions, background erythromycin resistance was undetectable, whereas 4 out of 5 isolates, including ATCC 33270 exhibited detectable levels of natural competence (Figure 2C).

**Figure 2.**
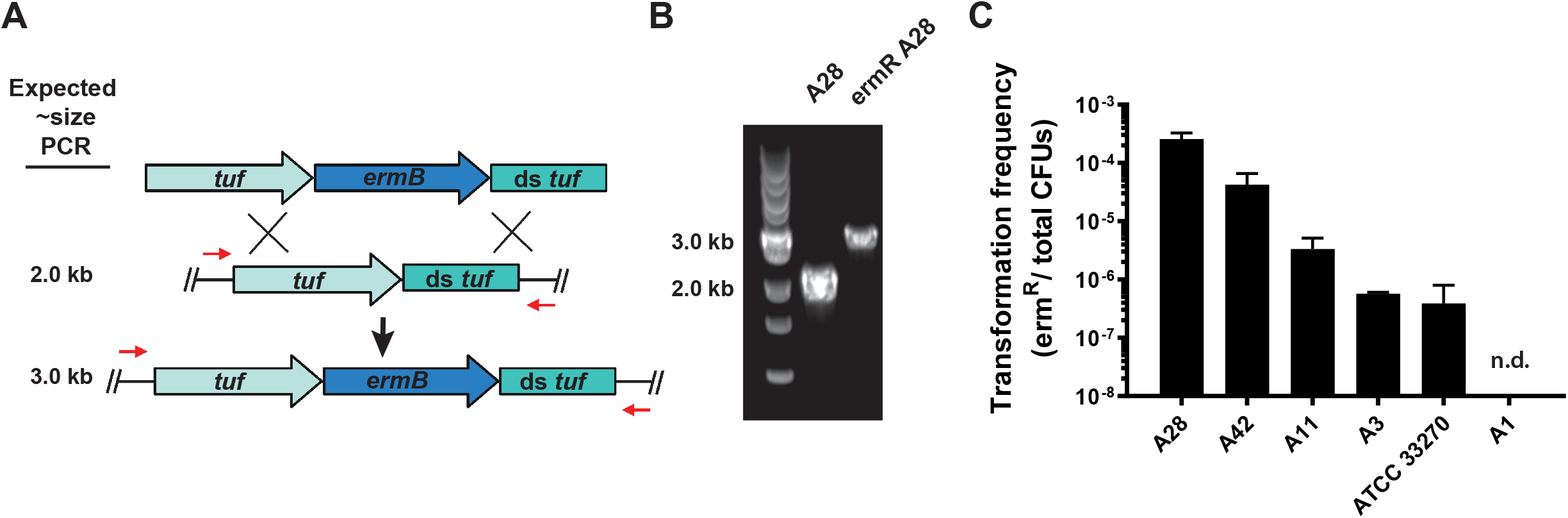
Insertion of resistance cassette into *P. micra* clinical isolates. (A) Mutagenesis construct (top) was inserted into the *P. micra* chromosome through homologous recombination (middle) yielding erythromycin resistant transformants (bottom). The primers used for PCR verification of genotypes are illustrated by red arrows. (B) Insertion of the erythromycin resistance cassette (~1 kb) was confirmed by PCR for a selected transformant (ermR A28). (C) Transformation frequencies of *P. micra* wild-type strains. Transformation frequencies are expressed as the ratio of erythromycin resistant CFUs to total CFUs. Values represent the averages from triplicate independent determinations ± SD. “n.d.” indicates a transformation efficiency below the detection limit of the assay.

### Comparison of construct homology vs. transformation efficiency

In an attempt to further bolster the transformation efficiencies of our isolates, we next tested a variety of mutagenesis constructs (Figure 3A) to examine the correlation between the sizes of the homologous DNA fragments and the resulting transformation efficiencies [26, 27]. Here, we focused on isolate A28, which displayed the highest level of natural competence using 1 kb homologous fragments (Figure 2C). No transformants were detected in the negative control samples or when using mutagenesis constructs containing homologous flanking regions of 250 bp (Figure 3B). An increase in the size of the homologous fragments from 1000 bp to 1750 bp resulted in >100-fold increase in transformation efficiency. A further increase in homologous fragment length from 1750 bp to 2500 bp only resulted in an additional ~15% increase in transformation efficiency, which suggested that transformation rates were approaching their maximum. Given the significant impact of larger homologous flanking regions upon the transformation rates of A28, we subsequently reexamined the transformation-negative strain A1 (Figure 2C) using 2.5 kb homologous flanking arms. Using this larger construct, it was then possible to detect reliable, albeit low levels of transformation in this strain (Figure 3B). These results demonstrate that the size of construct homology plays a significant role for natural transformation efficiency in *P. micra*, with homologous arms of ≥1750 bp being optimal for maximal rates of transformation.

**Figure 3.**
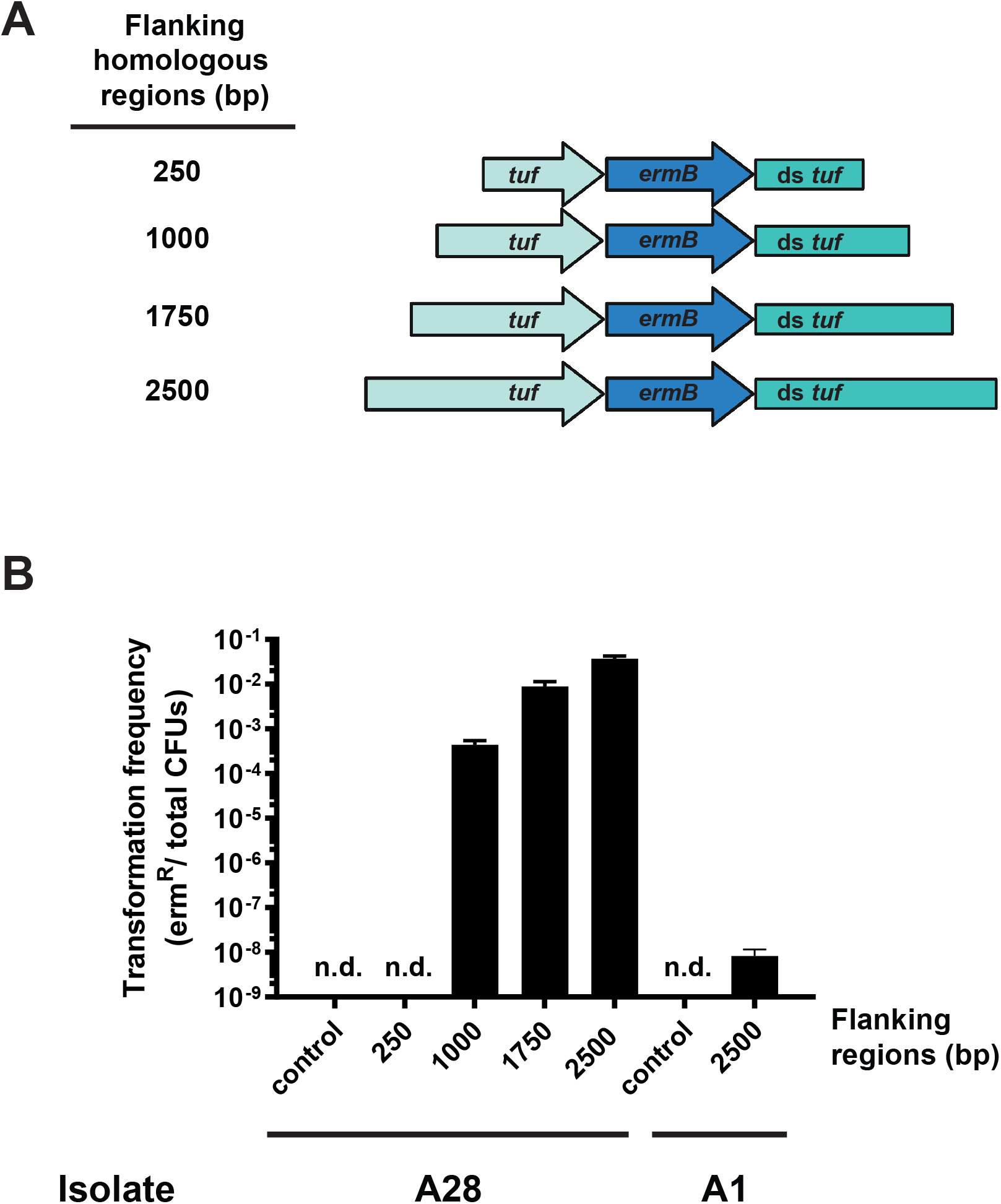
Comparison of homology vs. transformation efficiency in *P. micra*. (A) Mutagenesis constructs were generated with flanking regions of various sizes. (B) Wild-type clinical isolates A28 and A1 were transformed with the mutagenesis constructs illustrated in panel A. Values are averaged from triplicate independent determinations ± SD. “n.d.” indicates a transformation efficiency below the limit of detection of the assay.

### Targeted mutations of *recA* in *P. micra*

We were next interested to determine whether our *P. micra* transformation protocol could be used for targeted allelic replacement mutagenesis of *recA*. RecA is a nonessential DNA-dependent ATPase that plays a central role in homologous DNA recombination and repair [28, 29]. Given the requirement for homologous recombination during allelic replacement mutagenesis, it was anticipated that a *recA* mutant would exhibit a natural transformation defective phenotype, preventing subsequent mutagenesis using a marked linear DNA construct. Similar to other bacteria, we also anticipated a *recA* mutant to exhibit highly increased sensitivity to DNA damage. We tested this in strain A28 by first transforming a *recA* deletion construct comprised of an erythromycin resistance cassette (*ermB*) flanked by one kb DNA fragments homologous to the upstream and downstream regions surrounding *recA* (Figure 4A top). This resulted in a double crossover allelic exchange deletion mutant in which *recA* was replaced with *ermB* (Figure 4A middle and bottom). Since the transformation defects resulting from a *recA* mutation would likely preclude subsequent genetic complementation, it was necessary to first create an ectopically expressed *recA* knock-in mutant to serve as a recipient strain for a deletion of the native *recA* gene. A mutagenesis construct comprised of the *recA* ORF followed by a kanamycin resistance cassette (*aphAIII*) [25] was ligated to 2.5 kb segments of the *tuf* gene and its downstream sequence (Figure 4B top). After transforming this construct into strain A28, a double crossover event resulted in the insertion of an ectopic copy of the *recA* ORF and *aphAIII* directly downstream of *tuf* (Figure 4B middle and bottom). In this double *recA*-expressing background, we subsequently deleted the native *recA* gene using the same *recA* deletion construct described above (Figure 4A). Consequently, the resulting erythromycin/kanamycin resistant transformant harbored a deletion of the endogenous *recA*, while also expressing an ectopic copy of *recA* together with the *tuf* gene as part of an artificial operon. We subsequently compared the transformation efficiencies of these *recA* mutant strains using a PCR-generated amplicon of the rif^R^ *rpoB* gene previously generated in the strain ATCC 33270 (Figure 1B). The double *recA*-expressing strain (+A) and the complemented *recA* deletion strain (+A/ΔA) both exhibited similar transformation efficiencies to the wild-type (WT), whereas the *ΔrecA* deletion strain (ΔA) yielded no detectable levels of natural competence (Figure 4C). As an independent verification of the *recA* deletion phenotype, we further tested these strains for sensitivity to the genotoxic agent mitomycin C (MMC). As expected, the *ΔrecA* strain (ΔA) displayed a greater sensitivity to MMC at all of the tested dosages (0.5-4 μg mL^-1^), with a >3-log reduced survival rate observed at the highest MMC dosage (Figure 4D). In agreement with the transformation results, both the single and double *recA* strains behaved similarly to the wild-type. Taken together, these results demonstrate the successful implementation of a highly efficient *P. micra* mutagenesis protocol that can be employed for both targeted gene deletion and genetic complementation using naturally transformed PCR products.

**Figure 4.**
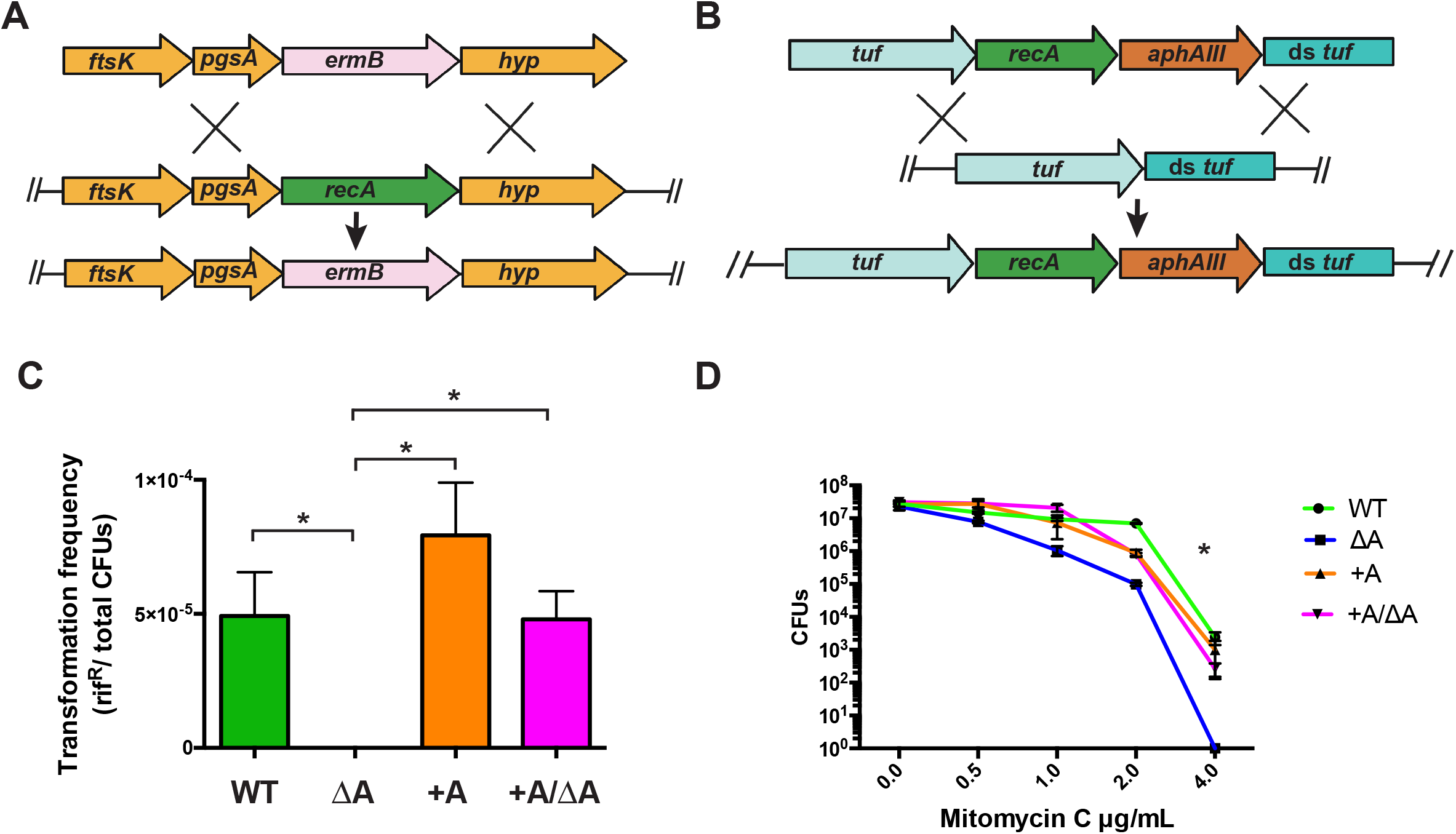
Targeted mutations of *recA* in *P. micra*. (A) The *recA* deletion construct (top) that was inserted into the *P. micra* A28 chromosome through homologous recombination (middle), resulting in allelic replacement of the *recA* ORF with an *ermB* cassette (bottom). (B) A *recA* knock-in mutagenesis construct (top) was inserted into the *P. micra* chromosome through homologous recombination (middle), resulting in the insertion of the *recA* ORF and *aphAIII* cassette downstream of EF-Tu (bottom). The Δ*recA* mutation was complemented by moving this mutation into the *recA* knock-in mutant background. (C) Transformation frequency of wild-type A28 (WT), A28 Δ*recA* mutant (ΔA), double *recA*-expressing knock-in mutant (+A), and complemented Δ*recA* mutant (+A/ΔA). Transformation frequency is expressed as the ratio of rifampicin resistant bacteria to total CFUs. (D) Mitomycin C survival curves. All values are averaged from triplicate independent determinations ± SD. For transformation assays, comparisons are indicated by brackets. For MMC experiments, comparisons are presented relative to ΔA. Statistical analysis was performed using a two-tailed Student’s *t*-test. “*” indicates a p <0.05.

### Creation of a renilla luciferase reporter

The high quantum yield and low background of luciferase-based reporter assays can provide a rapid and precise measure of gene expression over an exceptionally wide dynamic range. The superior signal-to-noise ratio of bioluminescent reporters has also supported noninvasive biophotonic imaging studies of mice orally infected with human oral streptococci [30]. To determine whether similar luciferase reporters would also function in *P. micra*, we developed a constitutive green renilla luciferase expression strain by inserting an independently translated green renilla (*renG*) luciferase gene and a kanamycin resistance cassette (*aphAIII*) immediately downstream of the *tuf* gene (Figure 5A). Green renilla luciferase (RenG) has been successfully employed in several species of oral *Streptococcus* and displays a much higher signal intensity compared to that of firefly luciferase [30]. Luciferase assays of this strain demonstrated extremely high levels of reporter activity that were >5-logs above the background readings of the unmodified parent strain A28 (Figure 5B). To gauge the detection limits and precision of the luciferase reporter readings in this strain, we next examined the correlation between population size and luciferase bioluminescence values. A dilution series of the reporter exhibited an extremely strong linear correlation with bioluminescence, with precise reporter values detectable in as few as 10^2^-10^3^ total CFU (Figure 5C). This suggests that luciferase reporter assays can be accurately performed within an exceptionally wide range of *P. micra* cell densities.

**Figure 5.**
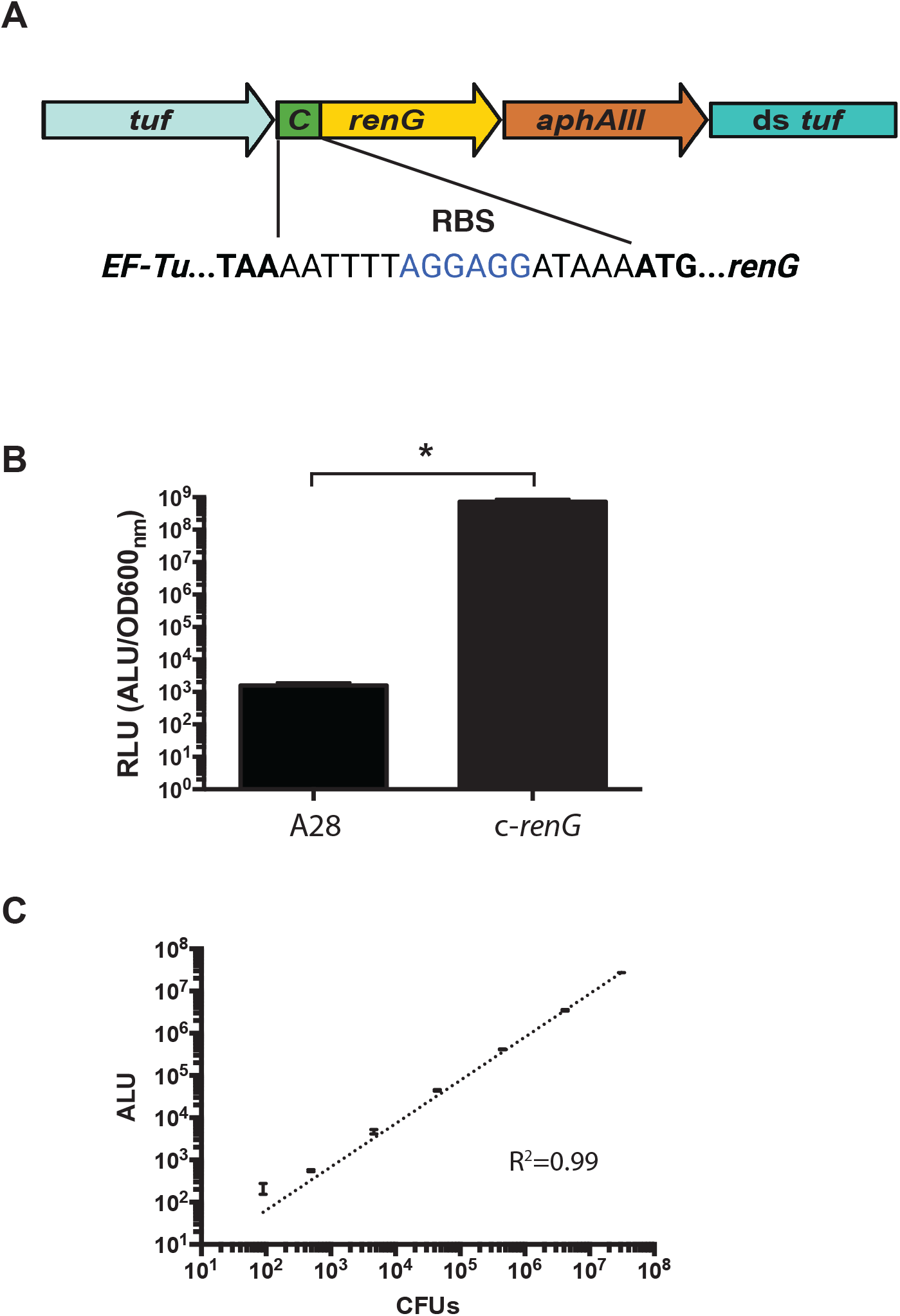
Green renilla luciferase (RenG) expression in *P. micra*. (A) Genomic context and sequence of a constitutively expressed *renG* transcription fusion to the *tuf* gene. (B) Luciferase activity was measured in wild-type strain A28 and the luciferase reporter strain (c-*renG*). Normalized luciferase activity (RLU) was calculated by dividing raw luciferase values by the culture optical density OD_600_ values. (C) Correlation between viable cell counts (CFUs) and luciferase reporter values in the constitutive luciferase reporter strain *c-renG* (R^2^ =0.99). All data points represent the average values ± SD from triplicate independent determinations ± SD. Statistical analysis was performed using a twotailed Student’s *t*-test. “*” indicates a p <0.05.

### Tunable-induction system in *P.micra*

The efficacy of the renilla luciferase reporter subsequently provided an avenue for developing an inducible expression system in *P. micra*. Regulated gene expression systems are powerful tools for genetic and functional studies. Optimal systems display a low basal expression along with strong and tunable levels of induction. We first tested the xylose-inducible expression cassette Xyl-S1 developed for oral *Streptococcus* species [31]. However, this only yielded nominal xylose-dependent regulation in *P. micra*. Therefore, we were curious whether a posttranscriptional regulatory system, such as those of ligand-inducible riboswitches [32], might perform better. To test this, we inserted the theophylline riboswitch [32] between the *renG* ORF and an ectopic copy of the constitutively-expressed *rpoB* promoter. This construct was then integrated onto the chromosome immediately downstream of the *tuf* gene transcription terminator to prevent transcriptional read-through from *tuf* (Fig. 6A). This strain was assayed for luciferase reporter activity over a range of theophylline concentrations, and we observed a titratable dose response curve with ~9-fold dynamic range (Figure 6C). This result is highly consistent with previous studies employing this riboswitch in other species [32]. Encouraged by the inducibility of this riboswitch in *P. micra*, we sought to improve upon its dynamic range by introducing a series of point mutations that were predicted to introduce a Rho-independent terminator-like element without triggering major changes to the overall riboswitch secondary structure (Figure 6B). Intriguingly, this modified reporter strain displayed a greatly enhanced response to theophylline, resulting in a >70-fold dynamic range (Figure 6C). Together, these results demonstrate the utility of *renG-encoded* luciferase as a sensitive reporter gene in *P. micra* as well as a riboswitch-based tunable gene expression system that exhibits low basal expression and robust inducibility.

**Figure 6.**
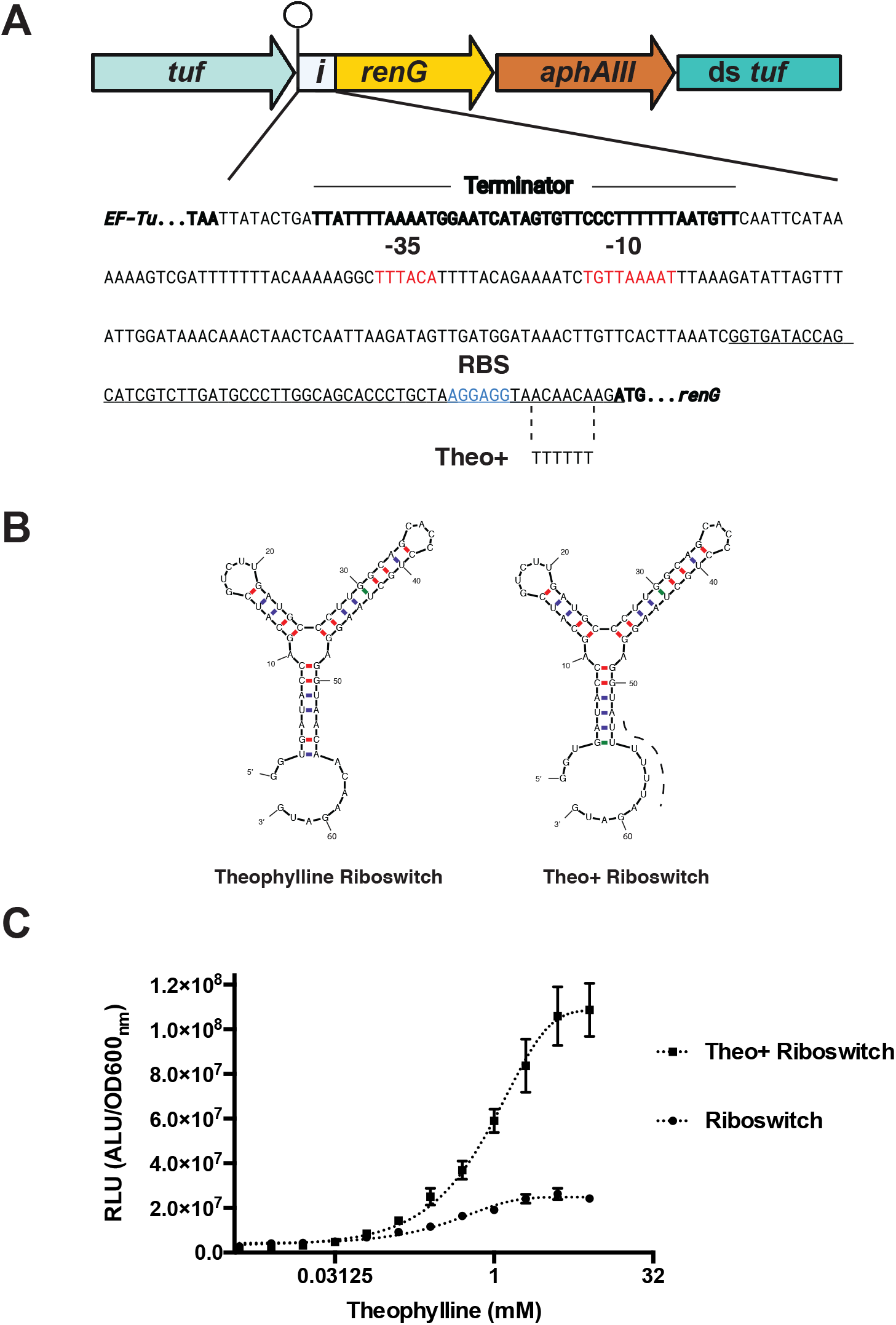
Regulated gene expression system in *P. micra*. (A) Genomic context (top) and corresponding sequence (bottom) for an inducible luciferase reporter strain (i-*renG*) in *P. micra*. The *tuf* gene Rho-independent terminator is indicated by a ball and stick icon. The sequences are shown for the promoter region of the luciferase reporter, with the original theophylline riboswitch (underlined) and Theo+ riboswitch (theo+*i-renG*) (dashed lines, bottom) indicated. Promoter elements are shown in red font, while the ribosomal binding site is in blue font. (B) The mFold webserver (http://www.unafold.org/mfold/applications/rna-folding-form.php) was used to predict the secondary structures of the original (left image) and Theo+ (right image) theophylline riboswitches. (C) The original and Theo+ riboswitches were compared for their abilities to regulate luciferase reporter activity over a range of theophylline concentrations. Normalized luciferase activity (RLU) was calculated by dividing raw luciferase values by the respective optical density OD_600_ values. Data points represent the average values ± SD from triplicate independent determinations ± SD.

### Development of *in vitro* transposon mutagenesis for Tn-seq studies

Transposon sequencing (Tn-seq) is a powerful forward genetic screening tool that has been successfully employed for genetic fitness studies of a number of pathogens [33] and pathobiont microbiome species [25, 34]. Given the high level of natural competence observed in *P. micra*, we were curious to determine the feasibility of implementing *in vitro* transposon mutagenesis using recombinant MarC9 transposase and the mini-Mariner transposon commonly used in Tn-seq studies (Figure 7A–B) [25, 33, 35]. Following *in vitro* transposon mutagenesis of *P. micra* gDNA, the reactions were transformed directly into the wild-type strain A28. With this approach, we readily obtained >6,000 transposon mutants/μg gDNA (Figure 7C and D). Considering that the *P. micra* genome is <2 Mb, a single *in vitro* mutagenesis reaction with several μg of *P. micra* gDNA would be sufficient to create densely saturated libraries of transposon insertions that are directly compatible with Tn-seq protocols. Overall, *P. micra* should be highly amenable to Tn-seq studies of its pathobiology.

**Figure 7.**
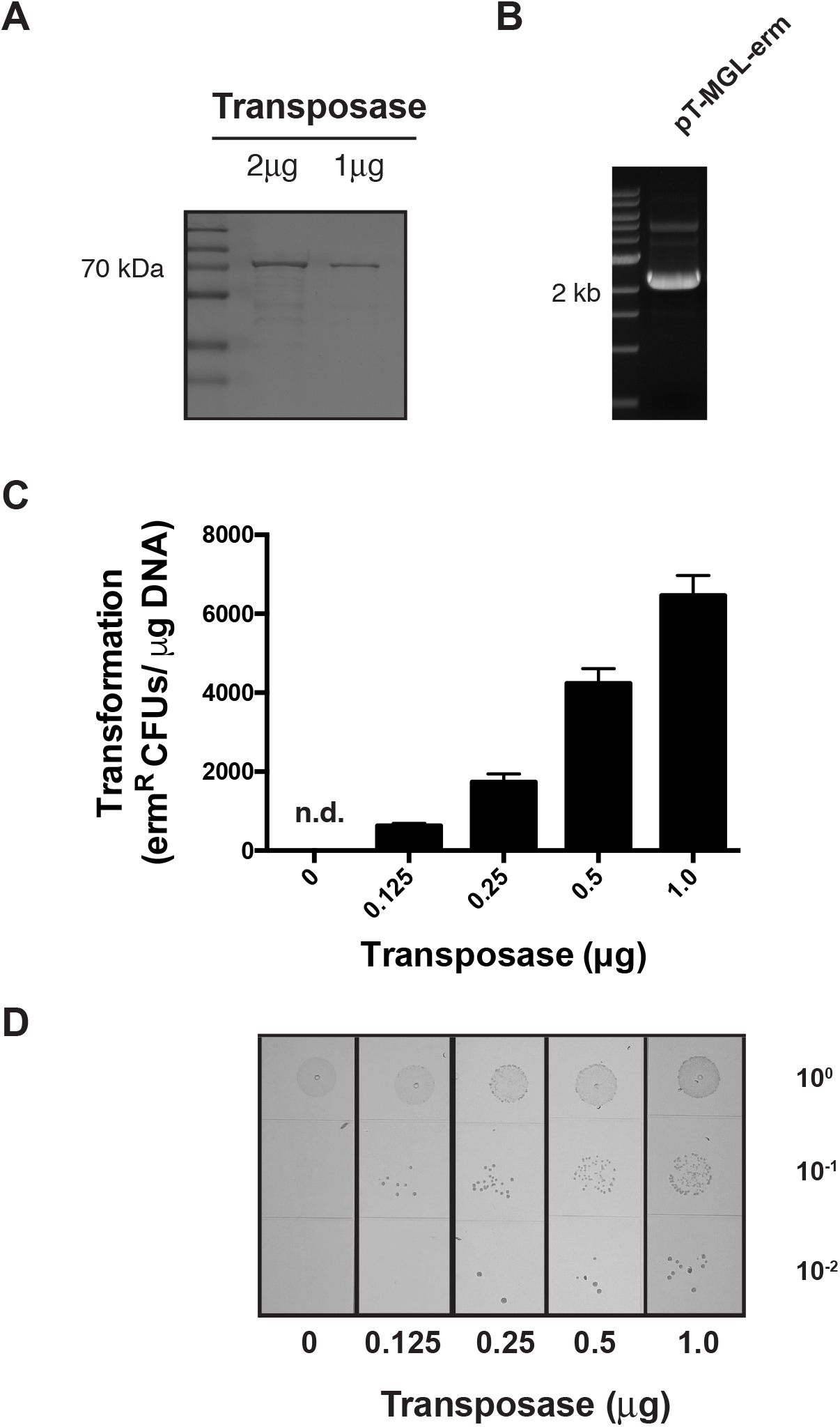
*In vitro* Mariner transposon mutagenesis in *P. micra*. (A) Purification of recombinant MarC9 transposase and (B) the mini-Mariner transposon plasmid pT-MGL-erm. (C) 1 *μg* of *P. micra* gDNA was mutagenized using 0.125, 0.25, 0.5, and 1.0 μg of recombinant MarC9 transposase with a fixed quantity of the mini-Mariner transposon. These reactions were subsequently transformed into wild-type *P. micra* strain A28. The transformation results are shown on the graph. Data are expressed as the average number of transposon mutants/μg of gDNA. Values are averaged from triplicate independent determinations ± SD. “n.d.” indicates a transformation efficiency below the detection limit of the assay.

## Discussion

*P. micra* is a notable pathobiont in the oral cavity due to its strong association with a variety of inflammatory oral diseases. *P. micra* is also commonly identified as a major constituent of acute and chronic infections at numerous other sites in the human body, including the abdomen, chest, spine, brain, blood, skin, and urogenital tract [36, 37]. Many *P. micra* infections have been described as polymicrobial, though a number of reports have implicated it as the sole source of infection as well [38, 39]. Interestingly, the presence of *P. micra* has also been strongly associated with multiple types of cancer. Numerous reports have identified a major enrichment of *P. micra* in colorectal tumors [5, 40–42], while others have reported an association of *P. micra* with oral and gastric cancers [4, 42, 43]. Given the substantial body of evidence for its role in human disease, the development of new genetic tools for *P. micra* is an essential step to reveal its unknown pathogenic mechanisms.

Here, we provide the first demonstration of *P. micra* natural competence. To the best of our knowledge, this is not only a first for the species, but it is also likely the first report of natural competence within the entire Tissierellia class, which is a subgroup in the Firmicutes phylum largely comprised of microbial dark matter. The use of natural competence for DNA transformation has numerous advantages over artificial methods, such as chemical treatments (e.g., CaCl_2_) and electroporation [44]. These approaches are also difficult to optimize for many fastidious anaerobes like *P. micra*, due to their sensitivity to excessive handling. Natural transformation can also be more efficient than those of artificial methods, as naturally transformed DNA enters the bacterial cytosol single-stranded [45]. Unlike duplex DNA introduced by artificial means, single-stranded DNA (ssDNA) is largely resistant to degradation by most bacterial restriction enzymes [46, 47]. Furthermore, ssDNA can serve as a more efficient substrate for homologous recombination compared to duplex DNA. The recombinase RecA is specifically activated by the presence of ssDNA to form pre-synaptic filaments, which in turn, promote RecA to scan for homologous sequences, induce synapse formation, and ultimately mediate recombination [48]. In our study, all of the *P. micra* isolates tested displayed some level of natural competence (Figure 2C and Figure 3B), though it is currently unclear why some strains exhibited higher transformation rates than others. A number of factors could contribute to these disparities such as strain-specific differences in the expression of competence related genes, variable responses to competence-specific environmental cues, and/or an altered expression of transformation-limiting bacterial factors (e.g. nucleases) [49]. Our results demonstrate that some of these limiting factors can be mitigated by increasing the lengths of the homologous fragments contained on the mutagenesis constructs (Figure 3B) [50]. Based on this, we speculate that endogenous exonuclease activity is likely to be a major determinant of the transformation efficiency observed with linear DNA constructs. However, it is also possible that longer segments of homology simply lead to higher rates of recombination.

After observing robust natural competence in *P. micra*, we examined the feasibility of creating *recA* deletion mutants. We chose *recA* as a test case due to its anticipated and easily measured phenotypes. As expected, the *recA* mutant exhibited a total loss of detectable natural transformability as well as a MMC-sensitive phenotype (Figure 4C and D). We were also able to complement these phenotypes by employing allelic exchange to insert the *recA* ORF immediately downstream of the constitutively expressed *tuf* gene, creating an artificial *tuf-recA* operon (Figure 4B). This further illustrates a straightforward approach for genetic complementation without requiring a shuttle vector.

The subsequent expression of luciferase in *P. micra* led us to consider its use as a reporter for the development of a regulated gene expression system. Previously, we developed a highly efficient xylose induction system (XylR/O) that functioned in several oral *Streptococcus* species [31]. Repressor/operator-based systems such as XylR/O (xylose) and LacR/O (IPTG) are contingent upon the uptake of inducer molecules as well as the heterologous expression of the corresponding repressor proteins. These can be of limited utility for bacterial species in which repressor protein expression or inducer permeability may be limiting factors. Indeed, our attempts to develop a XylR/O system in *P. micra* were largely unsuccessful. This led us to consider an alternative approach using a riboswitch-based regulatory system for posttranscriptional control of gene expression. Using a synthetic theophylline riboswitch [32], we demonstrated the potential utility of theophylline induction in *P. micra*, achieving up to a ~9-fold maximum induction of luciferase activity (Figure 6). In an attempt to further improve upon its performance, we introduced a series of riboswitch point mutations that were predicted to create a Rho-independent terminator-like structure in the absence of theophylline. Further studies would be required to confirm whether the mutant riboswitch truly functions in termination. Regardless, these mutations were able to further increase the dynamic range of the theophylline riboswitch by nearly an order of magnitude (Figure 6). This new version that we now refer to as the Theo+ riboswitch, yielded low basal expression, while providing robust and tunable inducibility. In addition to genetic complementation, this induction system is likely applicable for studies of both essential and toxic genes. It may also be useful for controlling target transcript levels and the timing of gene expression during infection studies.

Tn-seq is a powerful forward genetic screening tool used for genome-level assessments of genetic fitness. Based upon the efficacy of *P. micra* natural transformation, we predicted that it would be well suited for *in vitro* Mariner-based transposon mutagenesis. Considering that the Himar transposase specifically targets TA dinucleotides [35], the high A+T content of the *P. micra* genome (>70% A+T) makes it especially well-suited for Tn-seq analysis with Himar/Mariner. The availability of a high efficiency natural competence protocol also circumvents the requirement for complex *in vivo* transposition methods employing conjugation, temperature sensitive plasmids, and ectopically expressed transposases. Based upon the results shown in figures 7C and D, the current protocol should easily achieve the mutagenesis thresholds required for high-density Tn-seq analysis. Thus, when combined with the other genetic tools described in this study, one can now reliably perform nearly all of the molecular genetic approaches required to characterize *P. micra* pathobiology. As such, *P. micra* should also be added to the list of genetically tractable oral microbes.

## Materials and Methods

### Isolation of *Parvimonas* from Clinical Odontogenic Abscess Samples

Clinical sample collection and strain isolation protocols were reviewed by the Oregon Health and Science University Institutional Review Board (IRB) prior to the initiation of the study and deemed to be not human subject research. All clinical specimens were collected by clinicians in the OHSU Pediatric Dental clinic using materials generated during routine treatment procedures. Specimens were deidentified of all Protected Health Information (PHI) as part of the collection protocol. Clinical specimens were derived from a pediatric cohort undergoing tooth extraction due to odontogenic abscesses.

Briefly, abscess samples were collected using Dacron swabs and placed in pre-reduced transport media [21]. Abscess samples were vortexed and plated onto PMM agar media [12]. Plates were incubated in an anaerobic chamber (85% N_2_, 10% CO_2_, 5% H_2_) at 37°C for 4 to 7 days. Bacterial colonies producing a black precipitate were passaged on fresh PMM plates. Candidate isolates were imaged by microscopy for the expected cellular morphologies and then screened by PCR using the primers 16SFOR/REV. Sequence identification of 16S rRNA was confirmed using the expanded Human Oral Microbiome Database (www.homd.org) [51].

### Bacterial Strains and Growth Conditions

*Parvimonas micra* cultures were maintained in supplemented Brain Heart Infusion medium (sBHI), which consisted of base Brain Heart Infusion medium (BHI; Gibco, Gaithersburg, MD) containing 0.5% wt/vol yeast extract (Fisher, Fair Lawn, NJ), 0.005% wt/vol hemin (Sigma, St. Louis, MO), 0.001% wt/vol menadione (Sigma, St. Louis, MO), and 0.05% wt/vol cysteine (Sigma, St. Louis, MO). sBHI media and cultures were maintained in anaerobic conditions (85% N_2_, 10% CO_2_, 5% H_2_) at 37°C. For antibiotic selection, sBHI agar plates were supplemented with the following: rifampicin (5 mg L^-1^), kanamycin (300 mg L^-1^), and erythromycin (15 mg L^-1^).

To isolate spontaneous rifampicin resistant mutants, the *P. micra* reference strain ATCC 33270 was plated on sBHI agar containing rifampicin (5 mg L^-1^). A rifampicin resistant mutant (rif^R^) was screened by PCR using the primers rpoB-1240FOR and rpoB-2095REV. Sequence analysis of this conserved region of *rpoB* confirmed an A-to-T transversion resulting in an aspartic acid to valine substitution at amino acid 501 of RpoB.

### Electron Microscopy

Bacteria were grown overnight in sBHI on ITO coverslips and fixed in 2.5% glutaraldehyde, 2.5% formaldehyde in sodium cacodylate buffer (pH 7.4). Samples were rinsed in buffer, post-fixed in 2% osmium tetroxide for 30 minutes, rinsed in water, dehydrated in a graded ethanol series, and critical point dried in a Leica CPD300. Samples were mounted on aluminum stubs with a carbon tab and coated with 8nm carbon using a Leica ACE600 coater. Imaging was performed on a ThermoFisher Scientific Helios NanoLabG3 DualBeam scanning electron microscope.

### Extraction of genomic DNA from *P. micra*

Genomic DNA (gDNA) was purified using a phenol-chloroform extraction procedure [52]. Bacteria were harvested from sBHI agar plates, suspended in STES buffer (0.5M NaCl, 0.02 M EDTA, 0.2 M Tris-HCl (pH 8), 20 mg mL^-1^ lysozyme) and incubated at 55°C for 2 h. SDS was added to a final concentration of 1% followed by the addition of proteinase K (0.1 mg/mL) and incubation at 55°C for 1 h. One volume of phenol:chloroform:isoamyl alcohol (25:24:1) (Fisher, Fair Lawn, NJ) was added to the suspension and mixed gently. After centrifugation at 16,000 x g, the aqueous layer was collected and treated with RNase (50 μg mL^-1^) at 37°C for 2 h followed by a second extraction with phenol:chloroform:isoamyl alcohol. A 1:10 volume of 3 M sodium acetate (pH 5.5) and 3 volumes of ice-cold ethanol was added and incubated at −20°C for 24 h. DNA was pelleted by centrifugation at 16,000 x g, at 4°C for 30 min and subsequently washed with ice-cold ethanol (70%). The DNA pellet was dried in a laminar flow hood and resuspended in ddH_2_0.

### DNA transformation experiments

Transformation assays were performed similarly as previously described [24]. Briefly, *P. micra* was grown on sBHI agar for 72 h. Cells were harvested from plates and suspended into liquid sBHI and adjusted to an optical density OD_600_ of 0.4. Unless otherwise indicated, a total of 1 μg of gDNA (dissolved in 20 μL of ddH_2_0) was spotted on sBHI agar plates. After complete absorption of the gDNA into the agar, 20 μL of each bacteria suspension was pipetted over the DNA spots and absorbed. Plates were incubated in anaerobic conditions at 37°C for 24 h.

Following the incubation period, cells were then harvested into 50 μL of sBHI and vortexed. Serial dilutions were spread onto sBHI agar plates containing the appropriate antibiotic as well as on nonselective sBHI plates to determine total CFU counts. Transformation efficiency was calculated as the ratio of antibiotic resistant transformants to total CFU. For allelic replacement mutagenesis and genetic complementation, transformations were performed similarly as described above.

### *P. micra ermB* insertion mutagenesis constructs

Primers and mutagenesis constructs were designed using Serial Cloner software (https://serial-cloner.en.softonic.com). All primers used in this study are listed in Table S1. All PCR amplifications were performed using Phusion High-Fidelity Polymerase (Fisher, Fair Lawn, NJ). For insertion mutagenesis constructs, an erythromycin resistance cassette (*ermB*) was inserted directly downstream of the *tuf* gene. The *ermB* cassette was amplified using primers ermR-FOR/REV and the plasmid pJY4164 as a PCR template [25]. Upstream homologous regions were generated using the primer pair tuf-FOR/REV for all isolates. For downstream homologous regions, the primer pair down-tufFOR/REV was used with isolates A1, A3, and A28, while down-tufFORa/tuf-REV was used for strains A11 and ATCC 33270. gDNA from each respective *P. micra* strain was used as the PCR template. All amplicons were screened for size, column purified (Qiagen, Germantown, MD), and assembled using Gibson Assembly Master Mix (NEB, Ipswich, MA) as per manufacturer’s instructions. The assembled construct was amplified by PCR using the primer pair tuf-FOR/down-tuf-REV, screened for size, and column purified. Transformation assays using these constructs were performed as described above. Primer pair tuf-FOR*/down-tuf-REV* were used to screen select transformants for *ermB* insertion.

To generate *ermB* insertion constructs with flanking regions of varying sizes, gDNA from an *ermB* insertion mutant (ermR A28) was used as a template. The primer pairs tuf-250-FOR/REV, tuf-FOR/ down-tuf-REV, tuf-1750-FOR/REV, and tuf-2500-FOR/REV were used to generate homologous fragments of 250 bp, 1.0 kb, 1.75 kb, and 2.5 kb respectively. For transformation assays, a molar equivalent to 1 μg of the 1.0 kb homologous flank construct was used for all constructs. Transformation assays with these constructs were performed as described above.

### *P. micra recA* deletion and genetic complementation

For construction of the *recA* deletion mutant, an *ermB* cassette flanked by homologous upstream and downstream regions to *recA* was assembled. The *ermB* cassette was amplified using the primer pair ermR-FOR/REV and pJY4164 as the PCR template. Upstream and downstream homologous regions to *recA* were generated using the primer pairs up-recA-FOR/REV and down-recA-FOR/REV respectively, with strain A28 gDNA used as the PCR template. All amplicons were screened for size, column purified, and assembled using Gibson Assembly Master Mix. The assembled construct was amplified by PCR using the primer pair up-recA-FOR/down-recA-REV, screened for size, and column purified. Transformants were selected on agar plates supplemented with erythromycin.

To generate the *recA* knock-in strain, a construct was made containing the *recA* ORF along with a kanamycin resistance cassette *aphAIII* inserted immediately downstream of the *tuf* gene. PCR fragments homologous to *tuf* and its downstream sequence were generated using the primer pairs tuf-FOR/tuf-REVa and down-tuf-FORb/tuf-2500-REV respectively, with A28 gDNA serving as the PCR template. The *recA* gene was amplified using the primer pair recA-FOR/REV and A28 gDNA as the template. The kanamycin resistance cassette *aphAIII* was amplified using the primer pair kan-FOR/REV with plasmid pWVTK as the template [25]. All amplicons were screened for size, column purified, and assembled using Gibson Assembly Master Mix. The assembled construct was PCR amplified using the primer pair tuf-FOR/tuf-2500-REV and then column purified. Transformants were selected on agar plates supplemented with kanamycin. A complemented *ΔrecA* mutant strain was generated by transforming the *recA* knock-in strain with the *recA* deletion construct described above. Transformants were selected on agar plates supplemented with kanamycin and erythromycin.

To compare transformation efficiencies of the various *recA* mutants, PCR amplicons were generated with the primer pair rpoB-1240FOR/2095REV using gDNA from the spontaneous *P. micra* rifampicin resistant strain described above. PCR products were screened for size and column purified. Transformation assays were performed as described above using sBHI ± rifampicin. For mitomycin C (Sigma, St. Louis, MO) sensitivity experiments, each strain was suspended in sBHI at an OD_600_ of 1.0 and then serial dilutions were plated on sBHI agar plates containing varying dosages of MMC (0 μg mL^-1^, 0.5 μg mL^-1^,1 μg mL^-1^,2 μg mL^-1^, and 4 μg mL^-1^). Plates were incubated in an anaerobic chamber at 37°C for 72 h.

### Green renilla luciferase expression in *P. micra*

To express green renilla luciferase in *P. micra*, a construct was assembled containing the *renG* ORF along with a kanamycin resistance cassette *aphAIII* inserted immediately downstream of the *tuf* gene. PCR fragments homologous to *tuf* and its downstream sequence were generated using the primer pairs tuf-2500-FOR/tuf-REVa and down-tuf-FORb/tuf-2500-REV respectively, with A28 gDNA as the template. A PCR fragment of *renG* was generated using the primer pair renG-RBS-FOR/renG-REV with gDNA from strain brsRM-renG as the template [25]. The kanamycin resistance cassette *aphAIII* was amplified using the primer pair kan-FOR/REV and plasmid pWVTK as the template. All amplicons were screened for size, column purified, and assembled using Gibson Assembly Master Mix. The assembled construct was amplified by PCR using the primer pair tuf-2500-FOR/REV, screened for size, and column purified. The amplicon was transformed into wild-type *P. micra* strain A28 and the transformants selected on agar plates supplemented with kanamycin.

### Construction of *P. micra* inducible luciferase strain

For inducible expression of renilla luciferase in *P. micra*, a construct containing a *rpoB* promoter and riboswitch preceding *renG* followed by a kanamycin cassette was inserted directly downstream of the *tuf* gene rho-independent terminator. PCR fragments homologous to *tuf*, including its terminator and its downstream sequence were generated using the primer pairs tuf-2500-FOR/tuf-53-REV and down-tuf-FORb/tuf-2500-REV respectively, using strain A28 gDNA as the template. The riboswitch regulatory region was PCR amplified with the primer pair ribo-FOR/REV using a synthetic riboswitch DNA fragment (IDT, Coralville, Iowa) as the template (see Table S1). The *renG* gene was amplified with primer pair renG-FOR/REV using gDNA from the strain brsRM-renG. The kanamycin resistance cassette was amplified using the primer pair kan-FOR/REV and plasmid pWVTK as a template. All amplicons were screened for size, column purified, and assembled using Gibson Assembly Master Mix. The resulting construct was amplified by PCR using the primer pair tuf-2500-FOR/REV, screened for size, and column purified. This product was transformed into wild-type *P. micra* isolate A28 and transformants selected on agar plates supplemented with kanamycin.

For the Theo+ riboswitch variant, a construct was generated using the primer pairs tuf-2500-FOR/ theo-plus-ribo-REV and theo-plus-renG-FOR/tuf-2500REV using gDNA from the inducible mutant described above. PCR products were assembled using Gibson Assembly Master mix and amplified using the primer pair tuf-2500-FOR/REV. Transformants were selected on sBHI agar plates containing kanamycin.

### Luciferase assays

RenG expression strains were grown for 72 h on sBHI agar plates supplemented with kanamycin. Bacteria were suspended in liquid sBHI media with a range of theophylline concentrations (0 mM, 0.0078 mM, 0.0156 mM, 0.03125 mM, 0.0625 mM, 0.125 mM, 0.25 mM, 0.5 mM, 1 mM, 2 mM, 4mM, and 8mM) at an OD_600_ of 0.1 and then incubated in an anaerobic chamber at 37°C for 20 h. Coelenterazine-h solution (Prolume, Pinetop, AZ) was added to each sample (3.4 μg mL^-1^) and luciferase activity was measured on a Promega Glomax Discover luminometer. Optical densities were measured immediately after measuring luciferase activity for normalization. Normalized activity was expressed as relative light units (RLU), which is the luminescence value/OD_600_.

### Generation of a transposon insertion library in *P. micra*

A transposon library was generated in *P. micra* strain A28 using a previously described protocol [33]. Briefly, *in vitro* transposon mutagenesis was performed by combining 1 μg of gDNA from isolate A28, 1 μg of the Mariner transposon-containing vector pT-MGL-erm [25], and varying amounts of the MarC9 transposase (0 μg, 0.125 μg, 0.25 μg, 0.5 μg, and 1 μg). The mixture was incubated for 30°C for 1 h followed by a 75°C incubation for 10 min and then incubation on ice. The DNA was precipitated with ethanol and transposon junctions subsequently repaired. The resulting transposon reactions were transformed directly into *P. micra* isolate A28. Serial dilutions were plated on nonselective sBHI agar plates for total counts, while transposon mutants were selected on sBHI agar plates supplemented with erythromycin.

### Statistical analysis

All statistical analyses were performed using GraphPad Prism software to calculate significance via two-tailed Student’s *t-*test. Statistical significance was assessed using a cutoff value of P < 0.05.

## Acknowledgments

We are grateful to members of the Merritt laboratory for valuable insights regarding this manuscript. We thank Laura Higashi for help with scientific illustration. Images were prepared using Procreate, BioRender.com, Adobe Illustrator, and GraphPad Prism. Electron microscopy was performed at the Multiscale Microscopy Core, a member of the OHSU University Shared Resource Cores. This work was supported by NIH-NIDCR grants DE029612 and DE029492 awarded to J.K. and DE028252 awarded to J.M.

